# Hera-T: an efficient and accurate approach for quantifying gene abundances from 10X-Chromium data with high rates of non-exonic reads

**DOI:** 10.1101/530501

**Authors:** Thang Tran, Thao Truong, Hy Vuong, Son Pham

## Abstract

An important but rarely discussed phenomenon in single cell data generated by the 10X-Chromium protocol is that the fraction of non-exonic reads is very high. This number usually exceeds 30% of the total reads. Without aligning them to a complete genome reference, non-exonic reads can be erroneously aligned to the transcriptome reference with higher error rates. To tackle this problem, Cell Ranger chooses to firstly align reads against the whole genome, and at a later step, uses a genome annotation to select reads that align to the transcriptome. Despite its high running time and large memory consumption, Cell Ranger remains the most widely used tool to quantify 10XGenomics single cell RNA-Seq data for its accuracy.

In this work, we introduce Hera-T, a fast and accurate tool for estimating gene abundances in single cell data generated by the 10X-Chromium protocol. By devising a new strategy for aligning reads to both transcriptome and genome references, Hera-T reduces both running time and memory consumption from 10 to 100 folds while giving similar results compared to Cell Ranger’s. Hera-T also addresses some difficult splicing alignment scenarios that Cell Ranger fails to address, and therefore, obtains better accuracy compared to Cell Ranger. Excluding the reads in those scenarios, Hera-T and Cell Ranger results have correlation scores > 0.99.

For a single-cell data set with 49 million of reads, Cell Ranger took 3 hours (179 minutes) while Hera-T took 1.75 minutes; for another single-cell data set with 784 millions of reads, Cell Ranger took about 25 hours while Hera-T took 32 minutes. For those data sets, Cell Ranger completely used all 32 GB of memory while Hera-T consumed at most 8 GB. Hera-T package is available for download at: https://bioturing.com/product/hera-t

## 1 Introduction

In recent years, the emergence of single cell RNA sequencing technologies has allowed scientists to measure gene expression profile of thousands of individual cells simultaneously. The number of cells in each sequencing run has increased rapidly thanks to the invention of droplet-based protocols. In 10X-Chromium 3’ [1] protocol version 2, each transcript molecule is tagged with a 16-base-pair (bp) cellular barcode, and a 10-bp unique molecular identifier (UMI). Transcript molecules from the same cell are tagged with the same cellular barcode. UMIs are randomly generated, therefore different molecules from the same transcript within a cell have a very low chance of sharing the same UMI. This allows us to directly count the number of transcript molecules without PCR biases.

According to 10XGenomics specification [2], Cell Ranger uses STAR [3], a splicing-aware aligner, to align all reads against a reference genome and to get the best mapping loci for each read. Cell Ranger then uses a genome annotation to pick out reads that can be aligned to the transcriptome and discard the remaining reads.

Specifically, Cell Ranger uses the genome annotation GTF to bucket reads into *exonic, intronic*, and *intergenic*. A read is *exonic* if at least 50% of it intersects an exon, *intronic* if it is non-exonic and intersects an intron, and *intergenic* otherwise. Cell Ranger further aligns exonic reads against annotated transcripts, looking for compatibility. A read that is compatible with exons of an annotated transcript, and aligned with correct orientation, is considered mapped to the transcriptome. If the read is compatible with a single gene annotation, it is considered uniquely (confidently) mapped to the transcriptome. Only reads that are confidently mapped to the transcriptome are used for UMI counting [2].

While the description of Cell Ranger algorithm is highly reasonable, by analyzing its code and results, we identified the following limitations in its aligning procedure. Specifically, Cell Ranger (STAR) fails to correctly align reads in cases listed below:

- **Reads that span across small exons:** When reads span across multiple exons, they scatter into multiple fragments in the genome coordinate. We observe that when one of these fragments (exons) is small, Cell Ranger fails to detect correct alignments. As Cell Ranger (STAR aligner) maps reads to the genome using k-mers for seed identifications, it cannot detect seeds in small regions of the genome (corresponding to small exons). For these reads, Cell Ranger usually either fails to map, or maps them to another locus on the genome with a lower alignment score or cannot detect the exon splicing (reads aligned across exon boundary into intronic regions instead of skipping the intronic region) (Figure 1).
- **Splicing appears near read terminal:** Similar to the short exon case, when splicing positions are near a read terminal, the end fragment of the read cannot be hit by a seed. We observe that Cell Ranger also fails to align in this scenario and usually extends the alignment into the adjacent intron (Figure 2).
- **Errors (or SNPs) in reads conceal the correct anchor exons:** When mapping transcriptomic reads against a reference genome, SNPs or errors in reads can make it difficult to find the anchor seeds of reads on exons, especially short exons. This leads to incorrect alignment.
- **Incomplete report for equally aligned loci:** While using Cell Ranger on several data sets, we found some strange cases where reads that map perfectly to multiple positions on the genome (and also perfectly to the transcript sequences at these locations) are reported only once by Cell Ranger. This is a bug in the alignment tool (STAR) that Cell Ranger uses and we have filed a bug report to the authors of STAR aligner.

**Figure 1.**
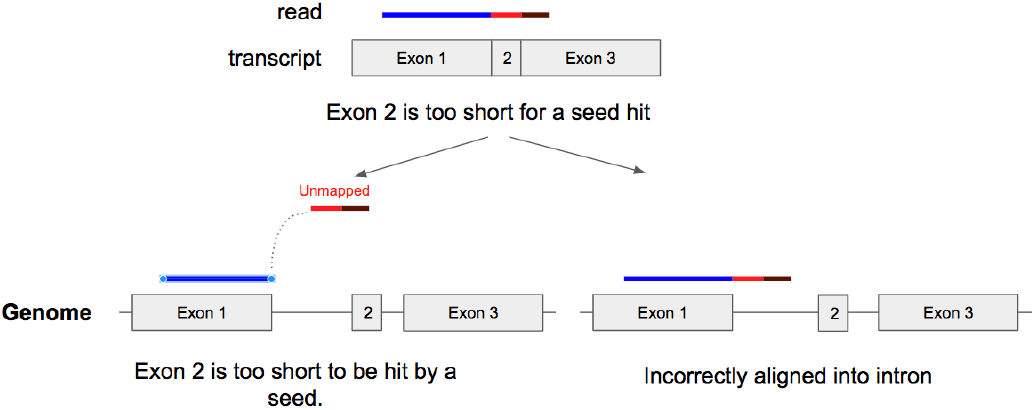
When a read spans across multiple exons, it is spliced into small fragments, making it difficult to align correctly on the genome coordinate.

**Figure 2.**
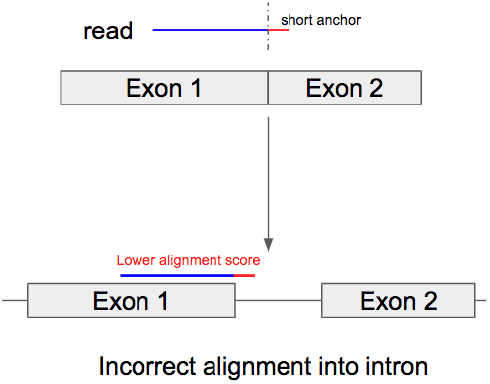
Splicing appears near a read terminal.

For example, in the 10X Chromium data set of 2k Brain Cells from an E18 Mouse [4], a read with ID @ST-K00126:491:HMV7GBBXX:2:2201:29812:36376 can be perfectly aligned to the following positions:

| Chr | Position | Cigar | Gene |
| --- | --- | --- | --- |
| 7 | 30484775 | 98M | None |
| 7 | 56014243 | 98M | None |
| 8 | 123206850 | 98M | None |
| 19 | 59322598 | 98M | ENSMUSG00000067038 |
| 10 | 23785587 | 91M224N7M | ENSMUSG00000061983 |

Yet Cell Ranger reports just one position.

| Chr | Position | Cigar | Gene |
| --- | --- | --- | --- |
| 10 | 23785587 | 91M224N7M | ENSMUSG00000061983 |

These incorrect alignment scenarios lead to the incorrect removal or selection of those reads in the Cell Ranger read selection procedure.

We introduce Hera-T, a new single-cell RNA-seq quantification algorithm for 10x-Chromium data. With many careful software engineering optimizations, Hera-T is 10 to 100 times faster than Cell Ranger, while consuming just a small memory footprint (smaller than 8 GB) for all benchmarked data sets.

Ignoring the difficult alignment cases that Cell Ranger failed to address, Hera-T and Cell Ranger produce almost identical results with the Spearman correlations larger than 0.99. As HeraT handles these difficult alignment scenarios correctly, we argue that it’s even more accurate than Cell Ranger.

For a single-cell data set with 49 million of reads, Cell Ranger took 3 hours (179 minutes) while Hera-T took 1.75 minutes; for another single-cell data set with 784 millions of reads, Cell Ranger took about 25 hours while Hera-T took 32 minutes. For those data sets, Cell Ranger completely used all 32 GB of memory while Hera-T consumed at most 8 GB.

## 2 Results

We benchmarked Hera-T against Cell Ranger [5] on 10X-genomics public single-cell data [6]. The code for reproducing the benchmarks is available at: https://github.com/bioturing/Hera-T-Benchmark. By the time we were finalizing this manuscript, Cell Ranger team released Cell Ranger 3. Therefore, we divided benchmark data into two groups:

- **v2 chemistry:** including 4 human and 7 mouse data sets.
- **v3 chemistry:** including 2 human and 4 mouse data sets.

To be consistent with the 10X-genomics public results, we ran Cell Ranger version 2.1.0 on v2 chemistry data sets and version 3.0.0 on v3 chemistry data sets. We benchmarked on a system with 56-core CPU and 32GB memory. Table 1 describes the running time and memory usage. As expected, there is an improvement in the performance of the new version vs the old version of Cell Ranger. For v2 chemistry data set, Hera-T is 50-100 times faster than Cell Ranger. While this number is about 10-48 times in v3 chemistry data set.

**Table 1.**
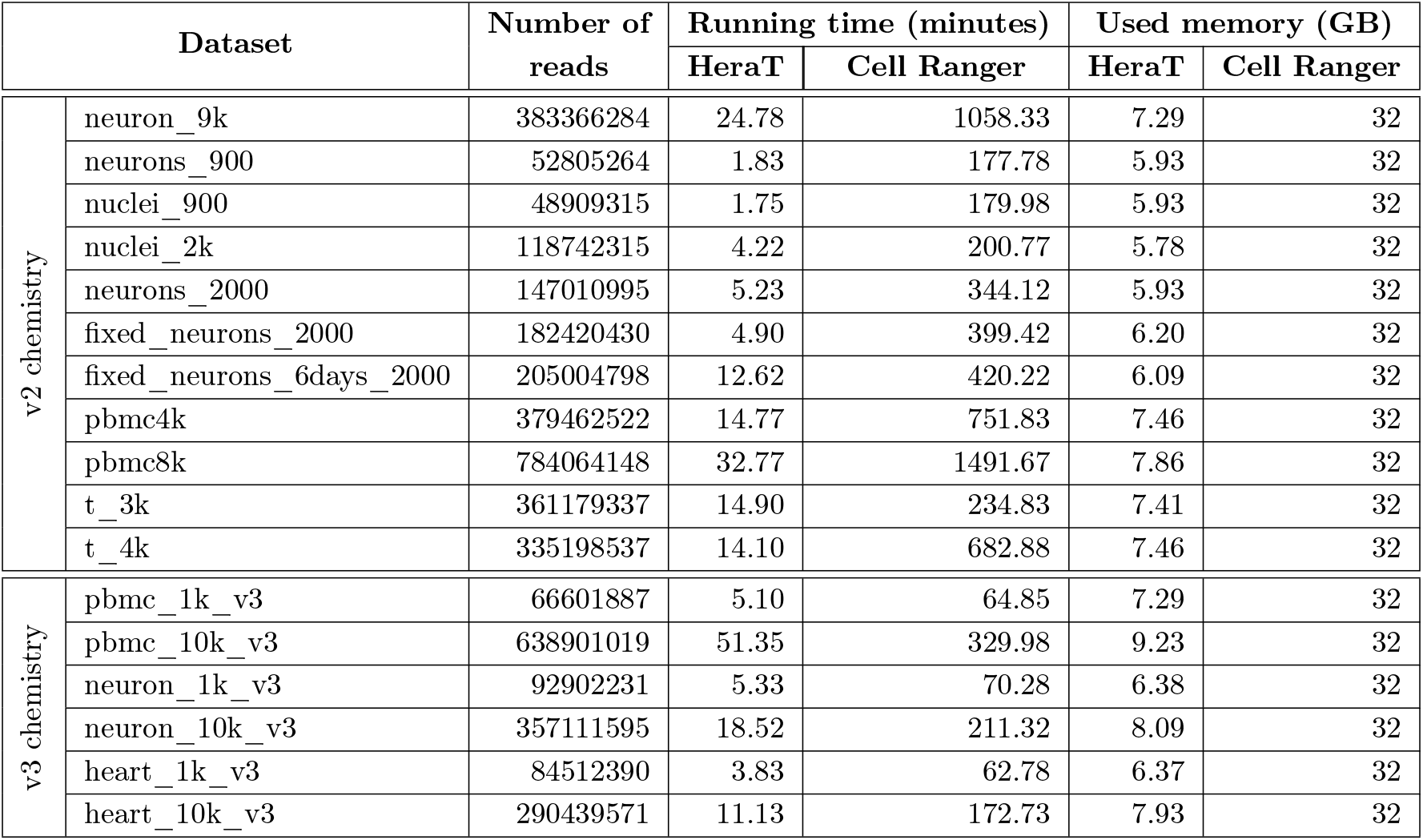
Memory and running time of Hera-T vs. Cell Ranger.

We use the following procedure to calculate the correlations between Cell Ranger and Hera-T results in order to assess their similarity.

- We get the set of shared barcodes reported by both tool (defined as set *W*).
- For each shared barcode, we filter out genes that have fewer than 2 UMI count in both tools (the set of remaining genes of *j*^th^ barcode is defined as *G_j_*).
- We compute the Spearman, Pearson, and expressed mean absolute relative difference (eMARD) scores between two vectors of UMI count (*x* and *y*). The eMARD score is calculated as equation 1 [7].

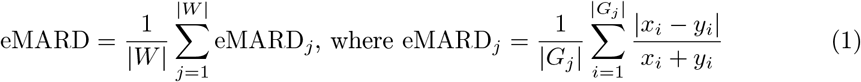
- Finally, we compare the mean and median scores of all shared barcodes between the two tools

Table 2 presents the correlation scores between Hera-T and Cell Ranger results.

**Table 2.** Correlation scores between **Hera-T** and **Cell Ranger** results

| Dataset |  | Spearman |  | Pearson |  | eMARD |  |
| --- | --- | --- | --- | --- | --- | --- | --- |
|  |  | Mean | Median | Mean | Median | Mean | Median |
| v2 chemistry | neuron_9k | 0.929576 | 0.933237 | 0.987605 | 0.988771 | 0.035957 | 0.034029 |
|  | neurons_900 | 0.933719 | 0.934995 | 0.986622 | 0.988992 | 0.038451 | 0.036891 |
|  | nuclei_900 | 0.922622 | 0.924616 | 0.984734 | 0.986303 | 0.042077 | 0.039727 |
|  | nuclei_2k | 0.769741 | 0.755276 | 0.994046 | 0.996602 | 0.086693 | 0.087956 |
|  | neurons_2000 | 0.919407 | 0.925676 | 0.993334 | 0.994163 | 0.050205 | 0.047587 |
|  | fixed_neurons_2000 | 0.921406 | 0.92555 | 0.990406 | 0.991377 | 0.050516 | 0.04905 |
|  | fixed_neurons_6days_2000 | 0.911156 | 0.914602 | 0.984244 | 0.985598 | 0.061773 | 0.060617 |
|  | pbmc4k | 0.972385 | 0.973166 | 0.999016 | 0.999438 | 0.028052 | 0.027688 |
|  | pbmc8k | 0.968846 | 0.969988 | 0.999111 | 0.999388 | 0.029999 | 0.02961 |
|  | t_3k | 0.950823 | 0.952669 | 0.99806 | 0.99887 | 0.045731 | 0.045149 |
|  | t_4k | 0.961283 | 0.963006 | 0.998421 | 0.999078 | 0.037364 | 0.036704 |
| v3 chemistry | pbmc_1k_v3 | 0.957775 | 0.962337 | 0.997647 | 0.999564 | 0.027869 | 0.025561 |
|  | pbmc_10k_v3 | 0.958123 | 0.960261 | 0.999319 | 0.999525 | 0.02869 | 0.027597 |
|  | neuron_1k_v3 | 0.955488 | 0.960985 | 0.994746 | 0.997093 | 0.022383 | 0.020621 |
|  | neuron_10k_v3 | 0.960629 | 0.963373 | 0.996117 | 0.996949 | 0.020031 | 0.01889 |
|  | heart_1k_v3 | 0.951225 | 0.962044 | 0.989389 | 0.993401 | 0.027405 | 0.023482 |
|  | heart_10k_v3 | 0.945173 | 0.958361 | 0.99146 | 0.992477 | 0.028488 | 0.023962 |

As mentioned above, Cell Ranger failed to align reads in some cases. We further removed those challenging reads together with reads having low alignment scores from the input data and re-benchmarked Hera-T vs. Cell Ranger. Specifically, we removed:

- Complicated spliced reads that Cell Ranger fails to map correctly. We identify those reads by comparing the alignments produced by Cell Ranger and bowtie2 on the transcriptome reference. When bowtie2 produces better alignment score compared to Cell Ranger, we consider that the read fails to be mapped by Cell Ranger.
- Reads can be mapped on multiple loci with equal alignment score but Cell Ranger only reports one of those.
- Reads with high error rates.

As a result, the two tools have almost identical results (Spearman and Pearson correlation are approximately 0.99). The results are presented in Table 3 and Figure 3. We also performed t-SNE to visualize these data, and included Alevin results into the picture (Fig. 5, 6). While the t-SNE plots of Hera-T and Cell Ranger results are almost identical, the plots from Alevin results are very different. The difference of Alevin can be tracked to two reasons. The first is that Alevin does not use genome reference for alignment, but only uses the transcriptome reference (for performance). The second source of the difference can be the way Alevin handles reads that map to mulitple locations, which is different from Cell Ranger and Hera-T.

**Figure 3.**
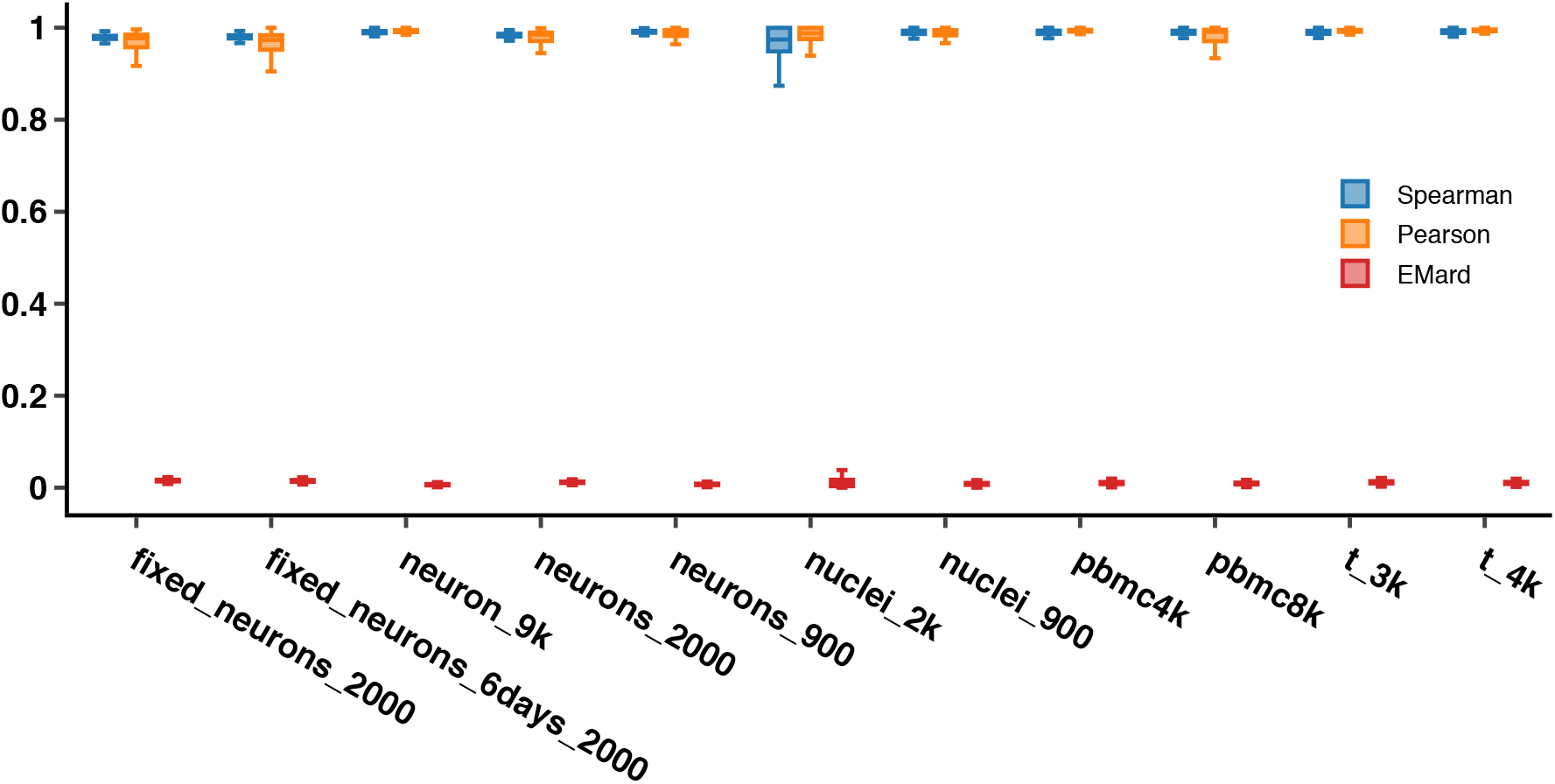
The distribution of Spearman, Pearson correlation scores, and eMard between Hera-T and Cell Ranger.

**Figure 4.**
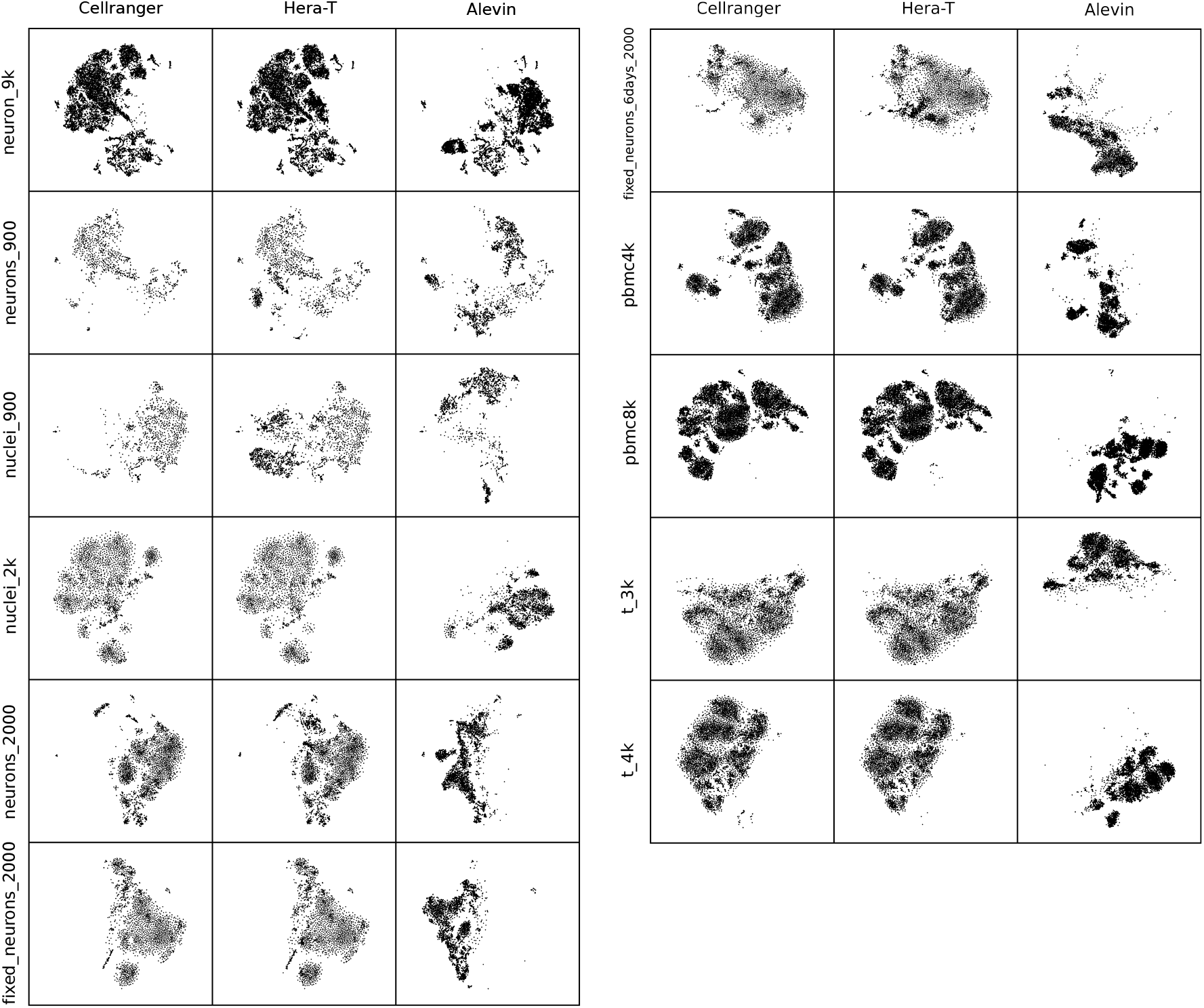
The t-SNE plots of Cell Ranger, Hera-T and Alevin results on v2 chemistry data. Cell Ranger and Hera-T plots are very similar, while Alevin’s plots are drastically different

**Figure 5.**
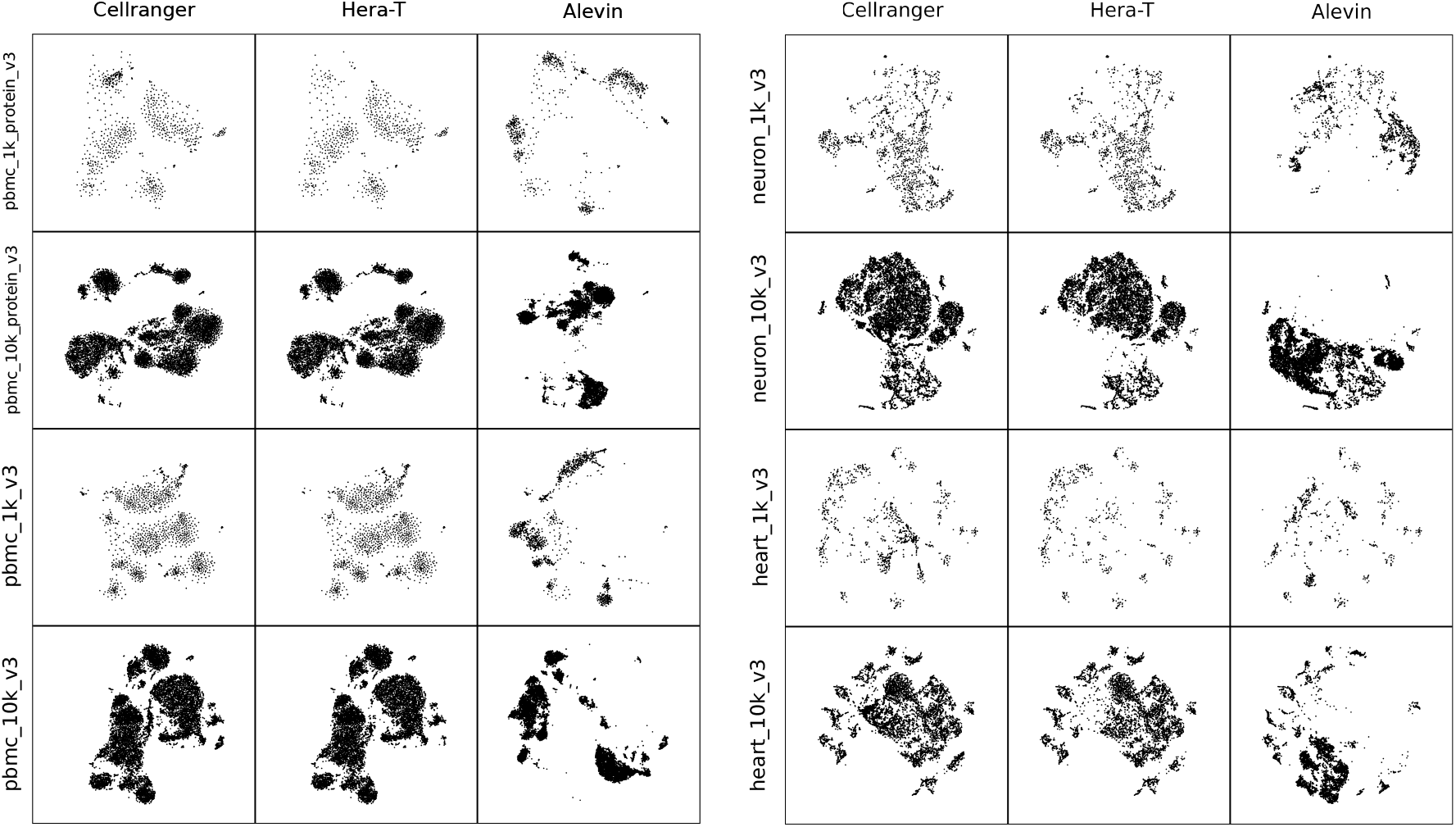
The t-SNE plots of Cell Ranger, Hera-T and Alevin results on v3 chemistry data. Cell Ranger and Hera-T plots are very similar, while Alevin’s plots are drastically different

**Table 3.** Correlation scores between **Hera-T** and **Cell Ranger** results after removing challenging reads

| Dataset | Spearman |  | Pearson |  | eMARD |  |
| --- | --- | --- | --- | --- | --- | --- |
|  | Mean | Median | Mean | Median | Mean | Median |
| neuron_9k | 0.9899 | 0.9907 | 0.9993 | 0.9995 | 0.0066 | 0.0066 |
| neurons_900 | 0.9909 | 0.9914 | 0.9994 | 0.9996 | 0.0073 | 0.0072 |
| nuclei_900 | 0.9890 | 0.9903 | 0.9992 | 0.9995 | 0.0084 | 0.0083 |
| nuclei_2k | 0.9687 | 0.9744 | 0.9989 | 0.9995 | 0.0118 | 0.0109 |
| neurons_2000 | 0.9830 | 0.9841 | 0.9993 | 0.9994 | 0.0118 | 0.0117 |
| fixed_neurons_2000 | 0.9787 | 0.9795 | 0.9989 | 0.9990 | 0.0154 | 0.0154 |
| fixed_neurons_6days_2000 | 0.9792 | 0.9806 | 0.9980 | 0.9984 | 0.0147 | 0.0148 |
| pbmc4k | 0.9898 | 0.9905 | 0.9997 | 0.9998 | 0.0102 | 0.0101 |
| pbmc8k | 0.9900 | 0.9905 | 0.9997 | 0.9998 | 0.0092 | 0.0091 |
| t_3k | 0.9897 | 0.9904 | 0.9997 | 0.9997 | 0.0118 | 0.0117 |
| t_4k | 0.9913 | 0.9920 | 0.9997 | 0.9998 | 0.0105 | 0.0103 |

## 3 Acknowledgements

This work was supported BioTuring Inc.

## References

[1] 10x-Genomics Single-Cell 3’-V2 Kit. https://teichlab.github.io/scg_lib_structs/data/CG000108_AssayConfiguration_SC3v2.pdf.

[2] Cell Ranger Algorithm Overview. https://support.10xgenomics.com/single-cell-gene-expression/software/pipelines/latest/algorithms/overview.

[3] Dobin, A. et al. Star: ultrafast universal rna-seq aligner. Bioinformatics 29, 15–21 (2013).

[4] 2k Brain Cells from an E18 Mouse. https://support.10xgenomics.com/single-cell-gene-expression/datasets/2.1.0/neurons2000.

[5] Zheng, G. X. et al. Massively parallel digital transcriptional profiling of single cells. Nature communications 8, 14049 (2017).

[6] 10x-Genomics Sinlge-cell sequencing data. https://support.10xgenomics.com/single-cell-gene-expression/datasets.

[7] Srivastava, A., Smith, T. S., Sudbery, I. & Patro, R. Alevin: An integrated method for dscrna-seq quantification. bioRxiv 335000 (2018).

